# Anti-citrullinated protein antibodies arise during affinity maturation of germline antibodies to carbamylated proteins in rheumatoid arthritis

**DOI:** 10.1101/2025.03.22.644346

**Authors:** Marta Escarra-Senmarti, Michael Chungyoun, Dylan Ferris, Jeffrey J. Gray, Felipe Andrade

## Abstract

Why autoantibodies in rheumatoid arthritis (RA) primarily target physiologically modified proteins, called citrullinated proteins, is unknown. Recognizing the inciting event in the production of anti-citrullinated protein antibodies (ACPAs) may shed light on the origin of RA. Here, we demonstrate that ACPAs originate from germline-encoded antibodies targeting a distinct but structurally similar modification, called carbamylation, which is pathogenic and environmentally driven. The transition from anti-carbamylated protein (anti-CarP) antibodies to ACPAs results from somatic hypermutations, indicating that the change in reactivity is acquired via antigen-driven affinity maturation. During this process, a single germline anti-CarP antibody transitions from anti-CarP to double positive (anti-CarP/ACPA) to ACPA according to the pattern and number of somatic hypermutations, explaining their coexistence and diverse specificity in RA. Artificial intelligence-based structural modeling revealed that an ACPA and its germline precursor exhibit distinct structural and biophysical properties, and pointed to heavy-chain tryptophan 48 (H-W48) as a critical residue in the differential recognition of citrullinated vs. carbamylated proteins. Indeed, a single methionine substitution in H-W48 changes the antibody specificity from ACPA to anti-CarP. These data indicate that the existence of germline-encoded anti-CarP antibodies is most likely the first event in the production of ACPAs during the early stages of RA development.

## Introduction

The production of antibodies to modified self-antigens is a hallmark in rheumatoid arthritis (RA), an autoimmune disease of unknown etiology characterized by immune-mediated damage to synovial joints ^1^. Antibodies to citrullinated and carbamylated proteins are of particular interest in RA because of their high prevalence, specificity, association with erosive disease, and appearance during the preclinical phase of RA ^2–6^. Citrullination is a physiological posttranslational modification (PTM) best known for its role in gene regulation and skin keratinization ^7^. During this process, arginine residues are converted to citrulline by the activity of the peptidylarginine deiminase (PAD) enzymes (**Fig. 1a**, upper panel) ^8^. Carbamylation is a chemical modification that results from the addition of a carbamoyl moiety to free functional groups of proteins, peptides, and free amino acids ^9–11^. This process is driven by environmental factors (such as smoke from biomass burning, biofuel usage, cooking, and tobacco smoking) and during inflammation, and mainly results from the reaction of isocyanic acid with a primary amine or a free sulfhydryl group ^2,9–11^. When carbamylation occurs on the ε-amino group of lysine residues, it generates homocitrulline residues (**Fig. 1a**, lower panel) ^9–12^.

**Fig. 1.**
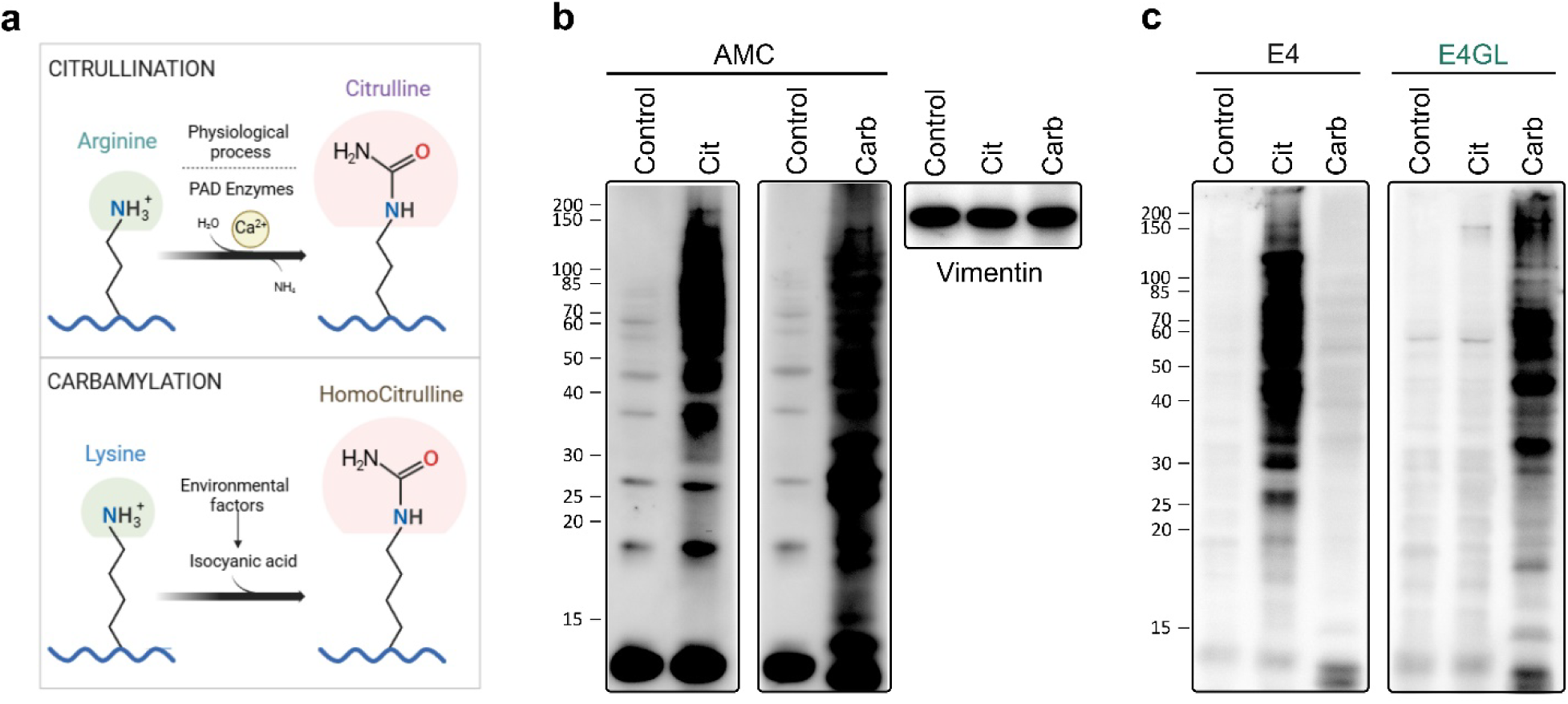
Monoclonal ACPA E4 changes reactivity to carbamylated proteins when reverted to its germline form. (**a**) Schematic view of the chemical structure of peptidyl-arginine and peptidyl-lysine, and their conversions to citrulline and homocitrulline residues (upper and lower panel, respectively). The amino acid side chain group that is modified is circled in green, and the resulting modification in pink. Citrulline and homocitrulline both contain a ureido group in the modified side chain (NH_2_CONH−). Homocitrulline is one methylene group longer than citrulline. PAD: Peptidylarginine deiminase. The figure was created with BioRender.com. (**b**) Mock or PAD2/PAD4 co-expressing 293T cells were stimulated with ionomycin to generate control and hypercitrullinated (Cit) cells, respectively. 293T cells were incubated with 1M potassium cyanate to induce carbamylation (Carb). Cellular citrullination and carbamylation were confirmed by AMC immunoblotting. Detection of vimentin was used as loading control (right panel). All immunoblotting assays used the same cell lysate batch and protein amount. (**c**) Lysates from control, Cit and Carb cells were tested by immunoblotting using 6.7 nM of antibody E4 or germline reverted E4 (E4GL). Data in **b** and **c** is representative from 2 and 4 independent experiments, respectively.

Citrullination and carbamylation have in common that they neutralize the positive charge of arginine and lysine, respectively, creating modified residues with similar structure, charge, and hydrophobicity (i.e., citrulline and homocitrulline, respectively). The only difference is that homocitrulline is one methylene group longer than citrulline **(Fig. 1a)** ^2^. Nevertheless, although citrullination and carbamylation generate nearly identical amino acid residues, these PTMs have significant differences in biology and potentially for RA ^2^. Citrullination is a physiological process across multiple tissues ^13^ and therefore, citrullinated proteins are poorly immunogenic *in vivo* ^14^. This is explained because, at least in mice, T and B cells reactive to citrullinated proteins are negatively selected during development ^15–17^. Indeed, the role of citrullinated proteins to induce arthritis in mice is controversial ^18–20^ and the antibody response is usually not citrulline-specific ^14,21,22^. In contrast, carbamylation is a pathological protein modification generated in conditions associated with oxidative stress ^12,23–26^. It is considered in the group of PTM derived products (PTMDPs), which includes other protein modifications such as oxidation, glycation, glycoxidation, lipoxidation, and carbonylation ^27,28^. PTMDPs are irreversible, accumulate throughout life, and are distinguished by the spontaneous binding to free reactive groups of proteins, which has a significant impact on protein structure and biological functions, especially in the context of aging, physiological stress, and inflammation ^24,27,28^.

Notably, different to citrullinated proteins, carbamylated proteins are highly immunogenic ^14,29^ and do not require a xenoreactive response to induce antibodies. Indeed, similar to other chemical modifications in proteins ^30^, anti-carbamylated protein (anti-CarP) antibodies are efficiently generated by immunization with autologous carbamylated proteins ^20,31,32^. Thus, although citrulline and homocitrulline are almost identical, the immune system can distinguish between the two PTMs under normal conditions. This is likely explained because different to citrullination, which is physiological, carbamylation is environmentally driven and thereby absent during B cell development. Thus, B cells able to react with carbamylated proteins are not subjected to negative selection. Interestingly, immunization with carbamylated peptides or autologous carbamylated albumin, but not with citrullinated peptides/albumin, renders mice susceptible to an aggressive form of erosive arthritis following a subsequent intra-articular injection of citrullinated peptides ^20^. Moreover, anti-CarP antibodies precede disease onset in monkeys with collagen-induced arthritis ^33^. Taken together, these data suggest a pathogenic link between carbamylation and citrullination in the induction of arthritis.

Although a potential mechanistic connection between anti-CarP antibodies and ACPAs remains unknown, the study of RA patient-derived monoclonal ACPAs supports that binding to carbamylated sequences results from cross-reactivity with ACPAs ^34^. Further understanding of the relationship between ACPAs and anti-CarP antibodies, however, has been hampered by the search for additional PTMs cross-reactive with ACPAs (e.g., acetyl-lysine and ornithine) ^35,36^, resulting in greater confusion about the origin of these antibodies ^2^.

Here, we provide a simple and rational explanation for the mechanistic link between citrullination and carbamylation in RA, integrating critical differences in their origin (physiologic vs. pathologic) and immunogenicity (tolerized vs. non-tolerized), which may explain why citrullinated proteins are abnormally targeted by the immune system in RA. Our studies indicate that antibody binding to carbamylated proteins is germline-encoded. B cells expressing germline anti-CarP antibodies escape negative selection, most likely because carbamylation is not physiologic, which explains why carbamylated proteins are normally immunogenic. However, since citrulline and homocitrulline are nearly identical, the data also indicate that germline-encoded anti-CarP antibodies can generate a mixture of anti-CarP, anti-CarP/ACPA and ACPAs as the result of antigen-driven affinity maturation, most likely in the context of genetic susceptibility, tobacco smoking, inflammation and hypercitrullination, among other factors. Overall, these findings imply that the presence of germline-encoded anti-CarP antibodies is the initial step in the production of disease-specific antibodies during disease initiation in RA.

## Results

### Somatic hypermutations dictate ACPA specificity for citrullinated or carbamylated cellular substrates

E4 is an RA patient-derived monoclonal ACPA that has been extensively characterized, including crystal structure analysis bound to citrullinated antigens and its effect on collagen antibody-induced arthritis (CAIA) in mice ^37–39^. While the specificity of E4 has been tested against synthetic citrullinated peptides ^38^, its reactivity to citrullinated and carbamylated cellular substrates has not been addressed. To gain insights into the reactivity of E4 to citrullinated and carbamylated cellular substrates and the role of somatic mutations in antigen binding, we generated recombinant antibodies in ExpiCHO cells using the E4 available sequence ^37,38^, as well as a version reverted to germline (both heavy and light chains). As substrates, 293T cells mock or transfected to co-express PAD2 and PAD4 were stimulated with ionomycin to generate control and hypercitrullinated cells, respectively. In addition, 293T cells were incubated with 1M potassium cyanate to generate carbamylated proteins (**Fig. 1b**). In contrast to lysates from control and carbamylated cells, E4 showed prominent binding to hypercitrullinated cells (**Fig. 1c**, left panel), demonstrating that citrullinated proteins are the major target of E4. When E4 was reverted to its germline form (E4GL), however, citrullination was no longer detected (**Fig. 1c**, right panel), indicating that reactivity to citrullinated proteins by E4 is gained as result of antigen-driven affinity maturation. Unexpectedly, while E4 showed poor recognition of carbamylated proteins (**Fig. 1c**, left panel), E4GL showed prominent binding to carbamylated proteins (**Fig. 1c**, right panel), implying that carbamylation is a primary target of E4GL. Moreover, the data indicate that, while somatic hypermutations promote E4 binding to citrullinated proteins, they have a significant impact on E4 targeting carbamylated proteins. Thus, unlike the current paradigm, these findings suggest that cross-reactivity plays a minimal role in the binding of ACPAs to citrullinated or carbamylated proteins. Instead, the data supports that the stage of antibody maturation seems to define the ACPA specificity to citrullinated or carbamylated proteins.

### The Chai-1 all-atom protein structure prediction method is capable of modeling the interactions of E4 with a citrullinated peptide

To understand the molecular basis that determines the distinct specificity between E4 and E4GL, we used structural modeling. Artificial intelligence (AI) methods for all-atom biomolecular structure prediction are capable of modeling the joint structure of complexes including proteins, nucleic acids, small molecules, ions, and modified residues ^40–44^. However, although these AI methods demonstrate substantially improved accuracy over many specialized tools for protein-ligand interactions, a comprehensive evaluation of their performance on specific antibody-antigen complexes involving non-canonical amino acids (NCAAs, for instance, citrulline and homocitrulline) remains unexplored. As no crystal structure of E4GL bound to a carbamylated peptide is available, but the structure of E4 bound to the citrullinated type II collagen peptide CII-Cit-13 (PDB: 5OCX) has been resolved ^38^, we first aimed to identify the AI model best suited for predicting the E4GL-carbamylated peptide complex. To achieve this, we benchmarked several all-atom biomolecular structure prediction methods, namely Chai-1 ^40^, AlphaFold3 ^41^, Boltz-1 ^44^, RoseTTAFold-AA ^42^, and Protenix ^43^.

Among the five structure prediction methods evaluated, only Chai-1, AlphaFold3, and Boltz-1 successfully positioned the citrulline residue of CII-Cit-13 within the paratope pocket of E4 (**Supplementary Fig. 1** and **Supplementary Table 1**). Both Chai-1 and AlphaFold3 predicted complexes and accurately depicted the three hydrogen bonds between E4 heavy (H) chain variable domain and citrulline that mediate its recognition. Two bonds involve the oxygen atom of the ureido moiety in citrulline, which interacts with the hydroxyl group of H-S51 and the amine nitrogen of H-W48, and a third bond between the ureido group nitrogen and the carboxamide oxygen of H-N105 ^38^. However, the Boltz-1 predicted complex lacked the crucial interaction between H-S51 and H-W48. Out of the eight regions of the E4 variable domain [i.e., six complementarity-determining region (CDR) loops, heavy chain (HC) framework, and light chain (LC) framework], Chai-1 outperformed AlphaFold3 in predicting the conformations of four regions: CDR H3, CDR L1, and both the HC and LC framework regions. Both methods performed equally well in predicting the conformation of CDR H1. Because Chai-1 accurately positioned citrulline within the E4 pocket and outperformed all other methods in predicting most of the E4 CDR loop and framework conformations—including the correct placement of H-W48, a crucial residue for citrulline binding in the HC framework ^38,39^—we chose this method to predict the conformation of E4GL with citrullinated and carbamylated peptides. Indeed, Chai-1 has been previously found to outperform AlphaFold3 at general protein-ligand complex prediction, and antibody-protein complex prediction ^40^.

### The differing sequence, structural, and biophysical properties of E4 and E4GL determine their respective antigen proclivity

To gain further insights into the role of somatic hypermutations in the transition from germline-encoded anti-CarP antibodies to ACPA, we investigated the differences in heavy-chain variable/light-chain variable (VH/VL) packing, charge, and solvent accessible surface area (SASA) when bound with either a citrullinated or carbamylated peptide. We predict the complex of E4 with CII-Cit-13 to represent an ACPA, and E4GL with a peptide sequence from carbamylated α1-antitrypsin (A1AT-HCit), which is targeted by anti-CarP antibodies and found in RA synovial fluid ^45^, to represent an anti-CarP antibody. We also predicted the complex of E4 with A1AT-HCit and E4GL with CII-Cit-13 to determine whether we observe the same binding preferences computationally as we did experimentally.

The VH/VL packing angle, distance, and opening angle of E4 and E4GL are similar in value (**Supplementary Table 2**), leading to a generally similar variable domain when overlapped with each other (**Fig. 2a**). However, E4GL displays a noticeably larger VH/VL opposite opening angle (**Supplementary Table 2**), which may explain the increased width of the antibody paratope pocket (**Fig. 2b**), as well as its preference for homocitrulline, which is one methylene group longer than citrulline (**Fig. 1a**). The CDR loops of E4 and E4GL antibodies differ in surface area, charge, and aromatic content. E4 generally has larger SASA values, particularly in H1, L1, and L2, indicating more exposed loops (**Supplementary Table 3**). E4 is more negatively charged in loops H1 and L1, while E4GL is often neutral or positively charged. E4GL also contains aromatic residues in loops like H2, where E4 lacks them (**Supplementary Table 3**). Despite these differences, both antibodies have similar charge and aromatic content in H3 (**Supplementary Table 3**). **Fig. 2c** further highlights the differences in charge distribution, with E4 exhibiting more negative charge on the HC side and the citrulline/homocitrulline (Cit/HCit) pocket, while E4GL has stronger positive charge areas on the LC side. The observed biophysical differences between E4 and E4GL, including variations in surface area, charge distribution, and loop structure, are key factors influencing their distinct antigen binding preferences. These results reinforce prior findings that somatic mutations in antibodies lead to significant biophysical changes compared to their germline versions, influencing their antigen binding properties and potentially driving their distinct specificity profiles ^46^.

**Fig. 2.**
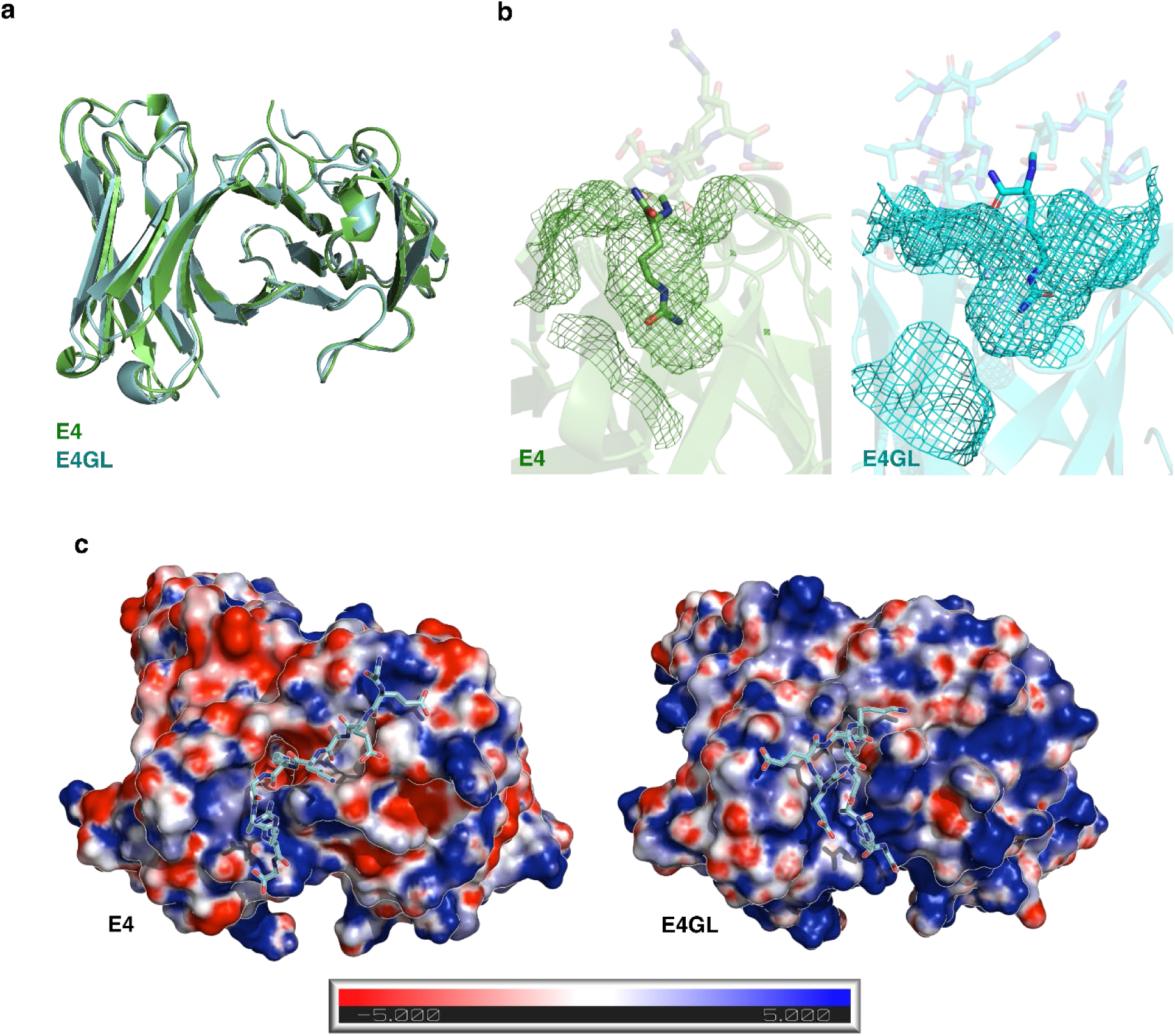
Biophysical characterization of deduced structures of E4 and E4GL. (**a**) Comparison of E4 and E4GL variable domains predicted by Chai-1. (**b**) A mesh representation of the E4 paratope with CII-Cit-13 and E4GL with A1AT-HCit reveals the variation in each antibody’s binding site architecture. (**c**) Charge-based surface models of E4 and E4GL. Protonation states calculated at physiological pH (7.4) with pdb2pqr ^66^; electrostatics calculated with APBS ^66^. The predicted CII-Cit-13 peptide is superimposed on the surface representation of each antibody to visualize approximate peptide placement relative to the charged patches.

### E4 and E4GL display opposing peptide binding characteristics

We initially predicted the structure of E4 and E4GL in complex with CII-Cit-13 or A1AT-HCit to exhaustively investigate the binding preferences of E4 and E4GL. The predicted complex of E4 with CII-Cit-13 shows key interactions between citrulline and three critical E4 HC residues ^38^ – H-W48, H-S51, and H-N105 (**Fig. 3a**). In contrast, the predicted E4-A1AT-HCit complex does not position homocitrulline within the antibody pocket (**Fig. 3b**). This difference in binding is also reflected in the predicted binding affinity, where E4 exhibits a stronger binding affinity to CII-Cit-13 than A1AT-HCit (**Supplementary Table 4**). While citrulline can still enter the E4GL pocket (**Fig. 3c**), structural differences between E4 and E4GL are apparent. In E4GL, the oxygen group of H-N105 points upward, whereas in E4 it points toward the center of the pocket. Additionally, two hydrogen bonds are lost in E4GL: the oxygen group of H-N105 no longer forms an intramolecular bond with the ureido group nitrogen, and H-W48 is too far from citrulline to interact (**Fig. 3c**). However, in the E4GL-A1AT-HCit complex, homocitrulline forms hydrogen bonds with H-N105 and H-S51 (**Fig. 3d**), with H-S51 oriented downward in both E4GL complexes, unlike the upward orientation observed in E4. The structural changes in E4GL shifts its predicted peptide affinity as well, where E4GL displays a stronger binding affinity to A1AT-HCit than to CII-Cit-13 (**Supplementary Table 4**). The structural differences between E4 and E4GL, particularly the orientation of H-N105 and H-S51 side chains, influence their peptide binding. Importantly, only H-W48 interacts with citrulline in E4, while in all other complexes, H-W48 is not involved. These differences in structure and interactions likely underlie the distinct binding preferences observed experimentally for E4 and E4GL, as well as the importance of H-W48 in citrulline but not homocitrulline binding.

**Fig. 3.**
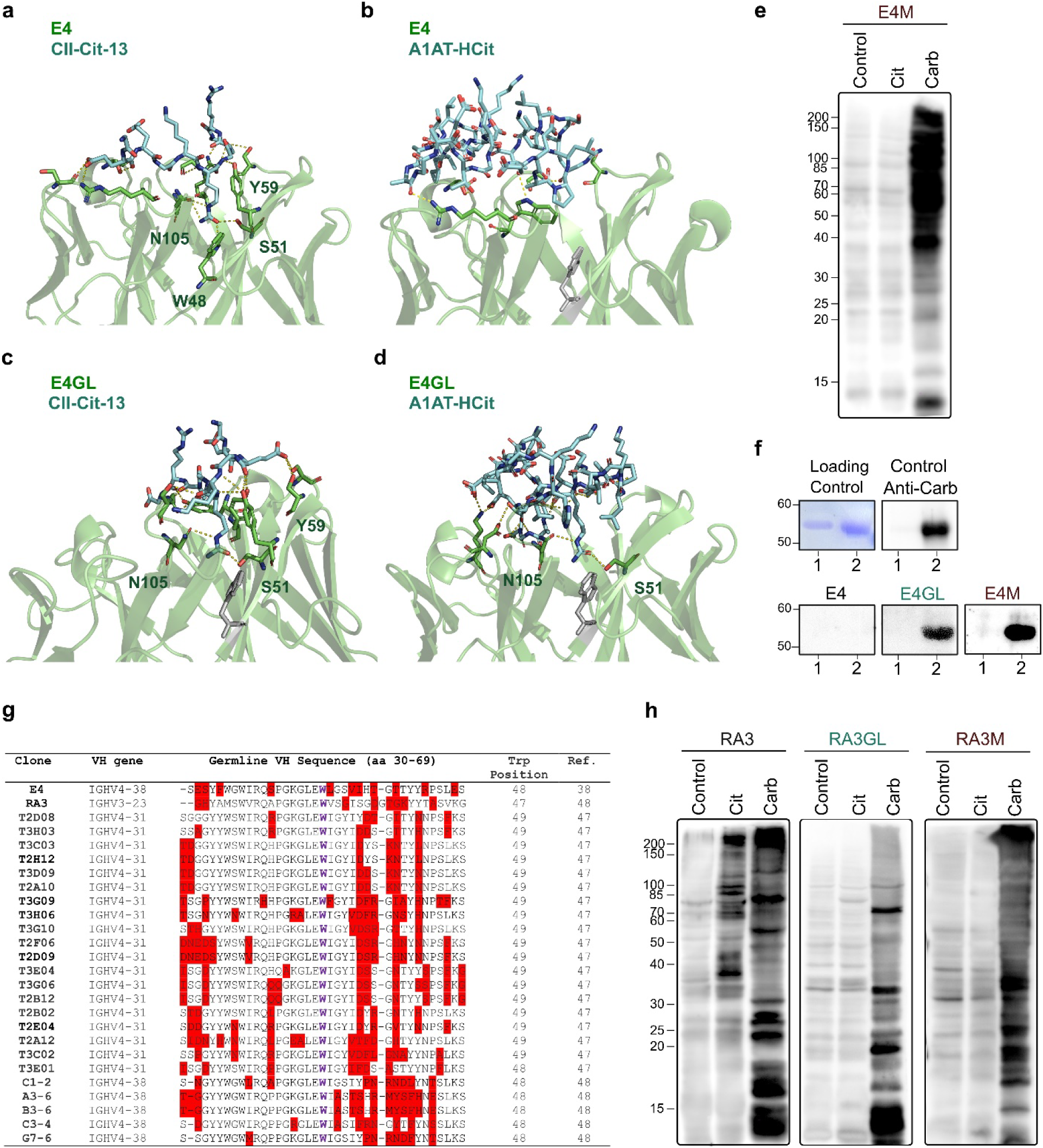
Heavy-chain tryptophan 48 (H-W48) dictates E4 binding to citrullinated or carbamylated proteins. Deduced structures from Chai-1 of (**a**) E4 bound to CII-Cit-13, (**b**) E4 bound to A1AT-HCit, (**c**) E4GL bound to CII-Cit-13, and (**d**) E4GL bound to A1AT-HCit. All antibody paratope residues (green) that interact with the bound peptide (cyan) are shown in stick representation. For clarity, only the antibody residues that interact directly with citrulline or homocitrulline are annotated. (**e**) Lysates from control, hypercitrullinated (Cit) and carbamylated (Carb) cells were tested by immunoblotting using antibody E4 in which H-W48 was changed to methionine (E4M). (**f**) Control (lane 1) and carbamylated (Carb, lane 2) α1-antitrypsin were detected by Coomassie staining (loading control) or by immunoblotting with a commercial antibody to anti-carbamyl-lysine (Cell Biolabs Inc., positive control) and antibodies E4, E4GL and E4M. (**g**) Sequence alignment of heavy chain (IgH) amino acid residues 33-66 from published RA patient-derived monoclonal ACPAs. Missense mutations are marked in red and conserved H-W47/48/49 in purple. Sequences were aligned using the multiple sequence alignment tool Clustal Omega from EMBL-EBI^67^. Except for monoclonal RA3M, which was used at 0.67 nM, all RA-derived monoclonal antibodies were tested at a concentration of 6.7 nM. Data in **e**, **f** and **h** is representative from 3 independent experiments.

### H-W48 is a key residue that dictates ACPA binding to citrulline or homocitrulline residues

While H-W48 is essential for citrulline binding by E4 (**Fig. 3a** and ^38,39^), it is worth noting that this residue is germline encoded in all human IGHV genes (except IGHV3-64 and IGHV3-64D) as either H-W47, H-W48 or H-W49. This implies that somatic mutations in residues other than H-W48 cause structural changes in E4 that positions H-W48 in the correct orientation to interact with citrulline. Indeed, when H-W48 was changed to methionine in E4 (E4M), the antibody lacked reactivity to citrullinated proteins (**Fig. 3e**), as previously described ^39^. However, it was surprising to find that, while E4M lacked its ACPA specificity, it regained reactivity to carbamylated proteins (**Fig. 3e**), mimicking the germline antibody E4GL (**Fig. 1c**, right panel). Moreover, consistent with the structural and experimental data, full-length carbamylated A1AT was only detected by E4GL and E4M, but not E4 (**Fig. 3f**). Collectively, the data imply that while H-W48 is essential for citrulline recognition by E4, this residue interferes with the efficient binding of E4 to homocitrulline. Thus, the specificity to citrulline or homocitrulline appears to be defined by H-W48, with its position determined by the pattern of somatic hypermutations. Notably, further analysis of published RA patient-derived monoclonal ACPAs ^37,38,47,48^ showed that although these antibodies are highly mutated, H-W47/48/49 is conserved (i.e., unmutated) among ACPAs (**Fig. 3g**), implying that like E4, the presence of tryptophan around position H-48 may also determine the specificity to citrulline or homocitrulline in other ACPAs.

To gain further insights into the origin of ACPAs, we produced three additional RA-derived monoclonals for which the VH and VL gene sequences are publicly available and have been shown to have strong reactivity to citrullinated peptides and some citrullinated proteins (RA3, A2-2 and F12-2) ^48^. Unexpectedly, except for a couple of proteins detected in hypercitrullinated cells, monoclonals A2-2 and F12-2 mainly recognized carbamylated proteins (**Supplementary Fig. 2a** and **2b**, respectively), suggesting that these are primarily anti-CarP antibodies. Nevertheless, when A2-2 and F12-2 were reverted to their germline forms (A2-2GL and F12-2GL), they remained anti-CarP, albeit with less reactivity (**Supplementary Fig. 2a** and **2b**, respectively), indicating that, like E4, they are derived from mutated germline antibodies to carbamylated proteins. In the case of RA3, it showed reactivity to both citrullinated and carbamylated proteins (Cit/CarP) (**Fig. 3h**, left panel) and consistent with E4, germline reverted RA3 (RA3GL) was only reactive to carbamylated proteins (**Fig. 3h**, middle panel) and a single mutation in H-W47 to methionine (RA3M) made the antibody anti-CarP specific (**Fig. 3h**, right panel).

### Single germline-encoded anti-CarP antibodies generate anti-CarP, ACPAs and double reactive anti-CarP/ACPAs as result of affinity maturation

Despite that RA-derived monoclonal antibodies catalogued as ACPAs (i.e., E4, RA3, A2-2 and F12-2) have distinct reactivity to citrullinated, carbamylated or both Cit/CarP, it is intriguing that, regardless of their specificity, they all derive from hypermutated germline precursors reactive to carbamylated proteins, implying a common mechanistic origin. To further understand the ontogenic relationship between anti-CarP, ACPAs and anti-CarP/ACPAs, we analyzed two sets of RA-derived monoclonal ACPAs that originate from single germline precursors. The first group includes four phylogenetically related ACPAs for which the VH and VL gene sequences are publicly available (termed T2H12, T2E04, T3G09 and T2D09) ^47^. In addition, we generated their common germline precursor (termed Gr1GL), including the putative germline-encoded CDR3 (**Fig. 4a**) ^47^. The second group includes five related monoclonal ACPAs (C3-4, G7-6, C1-2, A3-6 and B3-6)^48^, which also share a common germline precursor (termed Gr2GL) (**Supplementary Fig. 3**).

**Fig. 4.**
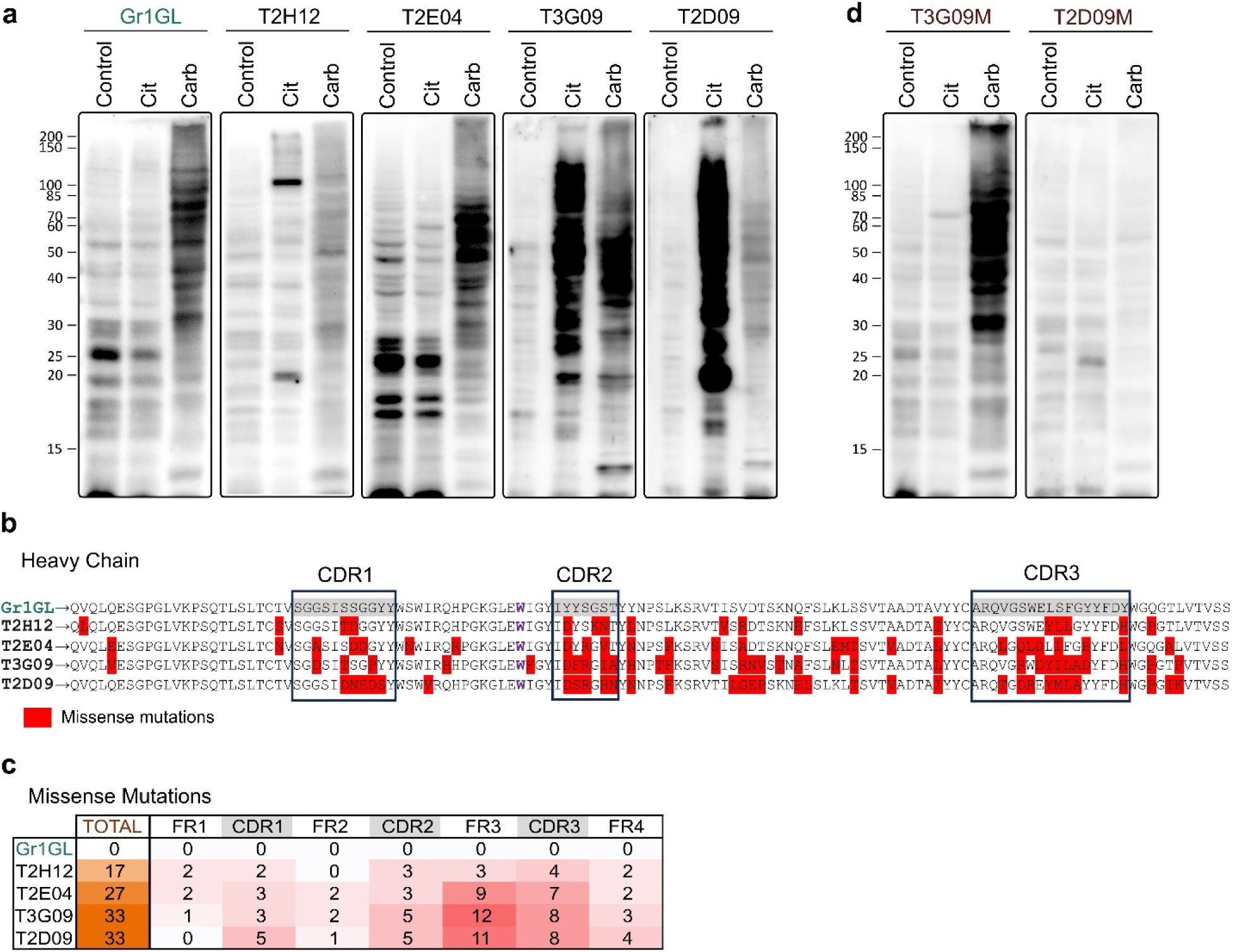
ACPAs, anti-CarP and double reactive anti-CarP/ACPAs comprise antibodies with distinct number and pattern of somatic hypermutations derived from single anti-CarP germline clones. (**a**) Lysates from control, hypercitrullinated (Cit) and carbamylated (Carb) cells were tested by immunoblotting using antibody Gr1GL, T2H12, T2E04, T3G09 or T2D09. (**b, c**) VH amino acid sequence alignment (**b**) and number of missense mutations (**c**) of antibodies shown in **a**. In **b**, missense mutations are marked in red and H-W49 in purple. FR, framework region. CDR, complementarity-determining region. (**d**) Lysates from control, Cit and Carb cells were tested by immunoblotting using antibody T3G09M or T2D09M. All RA-derived monoclonal antibodies were tested at a concentration of 6.7 nM. Data in **a** and **d** is representative from 3 independent experiments.

Consistent with the specificity of other germline reverted ACPAs (i.e., E4GL, A2-2GL, F12-2GL, and RA3GL), Gr1GL and Gr2GL only targeted carbamylated proteins but not control or hypercitrullinated cell lysates (**Fig. 4a** and **Supplementary Fig. 3a**), supporting the notion that ACPAs originate from germline-encoded anti-CarP antibody precursors. Interestingly, ACPAs derived from Gr1GL and Gr2GL showed different antigen specificities, generating antibodies with preferential binding to carbamylated proteins (e.g., T2E04), citrullinated proteins (e.g., T2D09 and C3-4), and with dual Cit/CarP reactivity (e.g., T2H12, T3G09, G7-6, C1-2, A3-6 and B3-6) (**Fig. 4a** and **Supplementary Fig. 3a**). Moreover, it is remarkable that the pattern of citrullinated and/or carbamylated proteins detected by each antibody is different, indicating that these antibodies have preference for distinct modified proteins, even when derived from the same germline precursor. The finding that somatic hypermutations in a single germline-encoded anti-CarP antibody can generate a mixture of anti-CarP, anti-CarP/ACPA and ACPA with distinct specificities explains their variable coexistence and diverse specificity among patients with RA.

Although Gr1GL-derived ACPAs that gained reactivity to citrullinated proteins (i.e., T3G09 and T2D09) had a higher number of missense mutations (**Fig. 4b** and **4c**), suggesting a transition from anti-CarP to ACPA according to the number of somatic hypermutations, this rule was not observed in Gr2GL-derived ACPAs, in which antibodies with a lower number of mutations (i.e., C3-4 and G7-6) (**Supplementary Fig. 3b** and **3c**) showed better reactivity to citrullinated proteins (**Supplementary Fig. 3a**). Thus, the data suggest that somatic hypermutations in germline-encoded anti-CarP antibodies can favor either specificity (ACPA, anti-CarP and/or anti-CarP/ACPA), which is likely determined by the type of antigen that dominates during the process of affinity maturation.

To further address the role of H-W48 (H-W49 in ACPAs derived from Gr1GL) in ACPA specificity, we mutated the tryptophan to methionine in three antibodies with the highest reactivity to citrullinated proteins (T3G09 to T3G09M, T2D09 to T2D09M, and G7-6 to G7-6M). Consistent with other ACPAs, antibodies T3G09M, T2D09M and G7-6M lacked reactivity to citrullinated proteins (**Fig. 4d** and **Supplementary Fig. 3d**), supporting that H-W48/49 is critical for binding to citrullinated proteins in all tested ACPAs. However, only antibodies T3G09M and G7-6M, not T2D09M, were strongly reactive to carbamylated proteins (**Fig. 4d** and **Supplementary Fig. 3d**), suggesting that in some ACPAs, other residues than H-W/47/48/49 are involved in their lack of reactivity to carbamylated proteins.

### Protein acetylation is a cross-reactive antigen for some ACPAs, but it is not a primary target of germline-encoded anti-CarP antibodies

Because acetylated-lysine residues resemble homocitrulline residues (except for the side-chain terminal amine, which is replaced by a methyl group) (**Fig. 5a**), and some ACPAs can be cross-reactive to acetylated proteins ^34,49^, it has been hypothesized that this modification may be relevant to the induction of ACPAs in RA. Using acetylated BSA (**Fig. 5b**), we found that all antibodies derived from E4 (E4GL, E4 and E4M) and RA3M showed reactivity to this antigen (**Fig. 5c** and **Fig. 5d**), indicating that different to homocitrulline and citrulline in proteins, acetylated-lysine residues are simply cross-reactive with no specificity driven by the E4 germline sequence, somatic hypermutations or H-W47/48/49. Notably, no other of the 12 ACPAs, including their germline and mutated H-W47/48/49 forms (except for RA3M), showed reactivity to acetylated BSA (**Fig. 5c** and **5d**). Thus, while acetylated-lysine residues can be cross-reactive to some ACPAs, which could be relevant for RA pathogenesis ^2^, our data do not support that this modification is involved in the origin of ACPAs in RA.

**Fig. 5.**
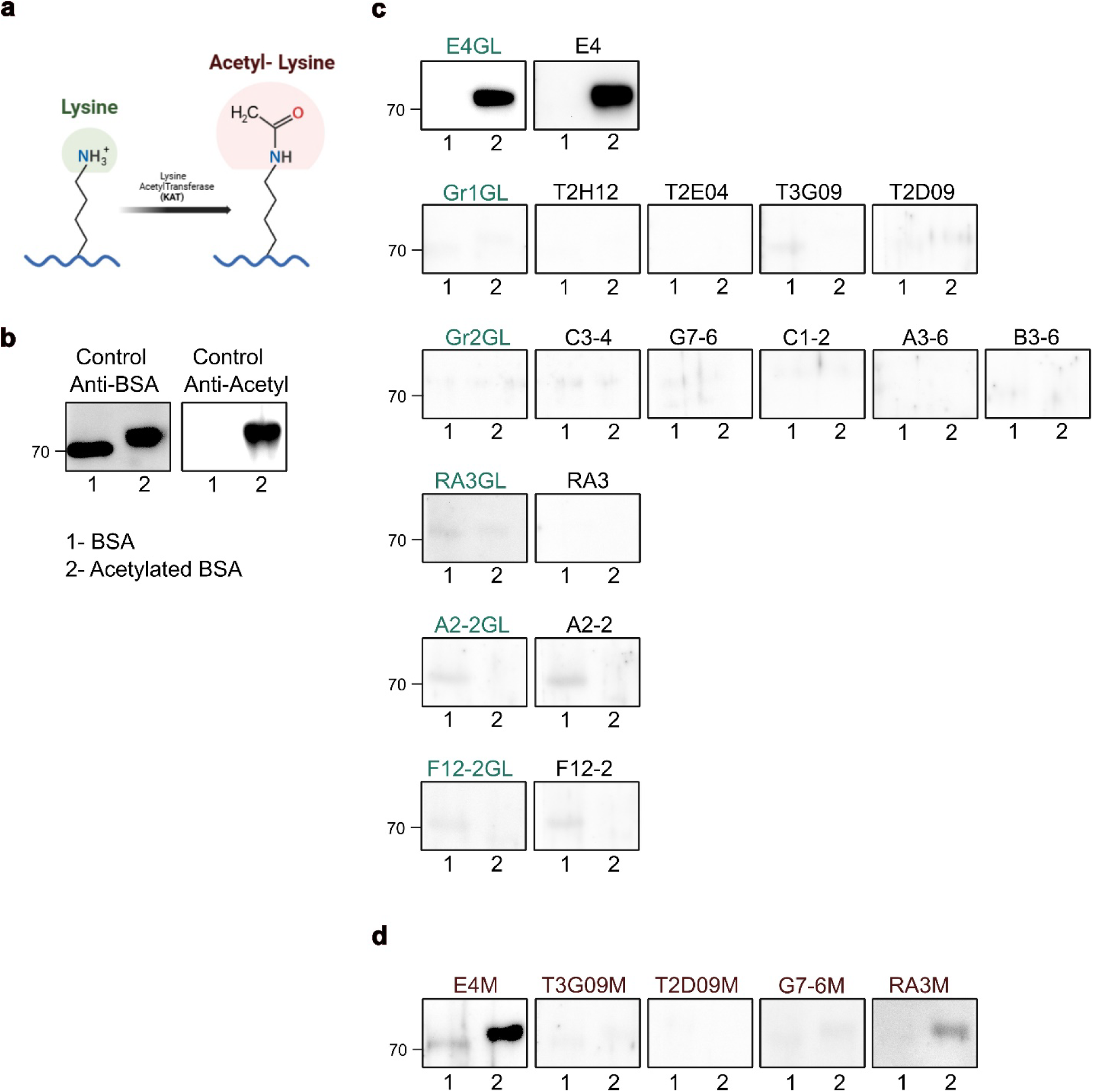
Protein acetylation is a cross-reactive target for certain ACPAs. (**a**) Schematic representation of the chemical conversion of peptidyl-lysine to an acetyl-lysine residue. The amino acid side chain group that is modified is circled in green, and the resulting modification in pink. Acetyl-lysine differs from homocitrulline by having a methylene group in place of the terminal amine found in the ureido group of homocitrulline. The figure was created with BioRender.com. (**b**) Bovine serum albumin (BSA) (lane 1) and acetylated BSA (lane 2) were detected using an in-house made anti-BSA mouse polyclonal antibody (loading control) or a commercial anti-acetylated-lysine antibody (Millipore Sigma) as the positive control. (**c, d**) Control and acetylated BSA were analyzed by immunoblotting using antibodies E4GL, E4, Gr1GL, T2H12, T2E04, T3G09, T2D09, Gr2GL, C3-4, G7-6, C1-2, A3-6, B3-6, RA3GL, RA3, A2-2GL, A2-2, F12-2GL and F12-2 **(c)** and H-M47/48/49 mutants **(d)**. Except for monoclonal RA3M, which was used at 0.67 nM, all RA-derived monoclonal antibodies were tested at a concentration of 6.7 nM. Data in **c** and **d** are representative of 2 independent experiments.

## Discussion

The analysis of serum samples collected years before clinical onset of RA has been critical in recognizing that the presence of disease-specific antibodies is one of the first events in the initiation of RA ^1,5,6^. Furthermore, the study of RA patient-derived monoclonal ACPAs has demonstrated that they are mostly generated from non-citrulline reactive precursors via antigen-driven affinity maturation ^50,51^. In this study, we showed that carbamylated proteins are the target of a set of ACPAs reverted to their germline form, implying that anti-CarP antibodies are precursors that can lead to the production of ACPAs during disease initiation in RA.

Depending on the pattern and number of somatic hypermutations, the data indicate that germline anti-CarP antibodies can generate distinct antibody subsets against carbamylated or citrullinated proteins, as well as double reactive antibodies. Because this combination of antibodies is most likely generated at random from different anti-CarP B cell precursors during the course of RA, it explains why studying the serologic appearance of anti-CarP antibodies and ACPAs during the preclinical and clinical phases of the disease has been insufficient to address the sequence of events leading to the origin of ACPAs. Instead, the analysis of autoantibodies in their germline form has been a useful strategy for understanding their antigenic origin and how somatic hypermutations create pathogenic antibodies, some of which exhibit multiple autoreactivities ^52,53^.

Interestingly, although autoreactive germline antibodies can be precursors of pathogenic autoantibodies ^52,53^, germline-encoded anti-CarP antibodies do not apply to this model because they are not properly autoreactive. Instead, they are directed to a novel protein structure, which is generated in the context of inflammation and noxious environmental stimuli ^2^. Thus, the presence of germline antibodies to carbamylated proteins is not explained by a defect in central tolerance. Rather, since carbamylated proteins are expected to be absent during normal development, B cells with immunoglobulin gene rearrangements reactive to carbamylated proteins would not be eliminated and would remain in the normal B cell repertoire in some healthy individuals.

Several RA-associated components are likely to concur to convert a non-pathogenic anti-CarP B cell into an ACPA producing cell. Chronic exposure to environmental factors that promote protein carbamylation, like tobacco smoking, may initiate the activation of naïve anti-CarP B cells ^54,55^. Similarly, the abnormal production of carbamylated proteins by leukocyte-derived peroxidases as result of chronic inflammation, such as at mucosal sites, is another potential source of carbamylated antigens ^25,56–58^. While these components can initiate an immune response to carbamylated proteins, this process is most likely self-limited, as it is with any foreign antigen under normal conditions. However, given that homocitrulline and citrulline are almost identical, the concurrence with hypercitrullination and RA-associated genetic susceptibility is likely to promote the further expansion and affinity maturation of anti-CarP B cells into ACPA B cells by targeting abnormally citrullinated proteins ^2^. For instance, microbial-induced hypercitrullination is an abundant source of citrullinated neoantigens ^59,60^, which may expose citrulline residues on structural epitopes recognized by anti-CarP antibodies.

Notably, this study demonstrates the power of AI-based all-atom protein structure prediction in modeling antibody-peptide interactions, particularly for antibodies with NCAAs. Our results show that Chai-1 effectively modeled the binding of E4 to citrullinated peptides, outperforming other tools, particularly in predicting CDR loop and framework conformations. Additionally, structural differences between E4 and E4GL, including changes in VH/VL packing, charge distribution, and aromatic content, explain their distinct binding preferences for citrullinated and carbamylated peptides. These findings underscore the importance of structural details like the orientation of key residues, such as H-W48, in determining antigen specificity.

Our work represents a significant step forward in using AI methods for modeling antibody-NCAA interactions, offering a novel approach for studying autoimmune diseases and therapeutic antibody design. By demonstrating the accuracy of Chai-1 in predicting antibody binding to modified peptides, we establish a foundation for more efficient antibody characterization in the absence of crystal structures. This study paves the way for future applications of AI-driven models in understanding antibody interactions with NCAAs. The success of our approach, which did not require a starting structure, highlights the potential of computational methods to predict complex biomolecular interactions and advance drug design and biomolecular research.

In summary, by combining experimental and AI-based structural data from RA-derived monoclonal ACPAs, our studies provide first evidence that RA most likely initiates from a repertoire of germline-encoded antibodies reactive to carbamylated proteins. In the context of RA susceptibility, chronic antigen stimulation and antigen-driven affinity maturation, these antibodies mature into a mixture of anti-CarP, ACPA and double reactive anti-CarP/ACPA, which are markers that precede the clinical onset of RA.

## Methods

### Cloning of monoclonal antibodies

The HC and LC variable gene sequences of monoclonal antibodies E4 (PDB: 5OD0_H, 50D0_L), T2H12 (GenBank: MH629725.1, MH629693.1), T2E04 (GenBank: MH629730.1, MH629692.1), T3G09 (GenBank: MH629731.1, MH629694.1) and T2D09 (GenBank: MH629717.1, MH629691.1), RA3 (GenBank: AJ430751.1, AJ430766.1), F12-2 (GenBank: AJ430750.1, AJ430769.1), A2-2 (GenBank: AJ430749.1, AJ430773.1), C3-4 (GenBank: AJ430740.1, AJ430759.10), G7-6 (GenBank: AJ430737.1, AJ430756.1), A3-6 (GenBank: AJ430743.1, AJ430762.1), B3-6 (GenBank: AJ430752.1, AJ430754.1) and C1-2 (GenBank: AJ430739.1, AJ430772.1) were synthesized through Integrated DNA Technologies (IDT), and cloned into expression vectors containing human Igγ1, Igκ or Igλ constant regions (kindly provided by Eric Meffre, Yale University School of Medicine, New Haven, CT).

### Reversion of SHM sequences to germline

The V(D)J germline sequences with the lowest number of mismatch nucleotides compared to mutated sequences were obtained using IMGT and IgBLAST, synthetized through IDT, and cloned into expression vectors containing human Igγ1, Igκ or Igλ constant regions. In Gr1GL, the putative germline VH and VL CDR3s were obtained from clade RA14 ^47^. In Gr2GL, the putative CDR3s were determined by the most conserved residues in antibodies C3-4, G7-6, C1-2, A3-6 and B3-6 in CDR H3 (ARGLSIYGTNDAFDV) and CDR L3 (AAWDDSLNGWV).

### Site-directed mutagenesis

A single-point mutation from tryptophan H-47/48/49 to methionine in antibodies E4, T3G09, T2D09, G7-6 and RA3 was performed using the Q5 Site-Directed Mutagenesis Kit from New England BioLabs. Mutations were confirmed by Sanger sequencing.

### Monoclonal antibody production

Monoclonal antibodies were produced using the ExpiCHO Expression System (Gibco) by co-transfecting plasmids encoding IgH and IgL according to the manufacturer’s instructions. Supernatants were collected on day 10 post-transfection, and antibodies were purified using Protein A beads (Pierce).

### Preparation of Citrullinated and Carbamylated HEK-293T Cell Lysates

HEK-293T cells were transiently transfected with mock or plasmids encoding PAD2 and PAD4 using Lipofectamine 2000 (Invitrogen). After 48 hours, the cells were incubated with 5 µM ionomycin and 2 mM calcium in HBSS supplemented with 10mM HEPES for 2 hours at 37°C. Following incubation, the cells were pelleted, washed twice with 1× PBS, resuspended in SDS sample buffer, and boiled. For carbamylation, HEK-293T cells were resuspended in 1 M potassium cyanate and sonicated in the presence of 1% Halt Protease Inhibitor Cocktail (Thermo Fisher). After overnight incubation at 37°C, the sample was extensively dialyzed against 1× PBS. Citrullination and carbamylation were confirmed by anti-modified citrulline (AMC) immunoblotting_61._

### Immunoblotting

Control, citrullinated and carbamylated cell lysates, as well as bovine serum albumin (BSA) (ThermoFisher), acetylated BSA, alpha-1-antitrypsin (A1AT), and carbamylated A1AT (Cayman Chemical) were analyzed by electrophoresis on 12% (lysates) or 10% (purified proteins) SDS-polyacrylamide gels, followed by protein transfer to nitrocellulose membranes for immunoblotting. Except for monoclonal RA3M, which was used at 0.67 nM, all RA-derived monoclonal antibodies were tested at a concentration of 6.7 nM. Images were acquired using a ProteinSimple Fluorochem-M digital imager.

### Biomolecular complex prediction

This section outlines the methods used for predicting biomolecular complexes based on given sequence representations. All biomolecular structure prediction methods investigated utilize a sequence representation of the wildtype or germline antibody sequence, along with a Simplified Molecular Input Line Entry System (SMILES) string of the citrullinated or carbamylated peptide. The peptide SMILES strings are generated using PepSMI ^62^.

#### Wildtype heavy chain

QVQLEESGPGLVRPSETLSLSCTVSGFPMSESYFWGWIRQSPGKGLEWLGSVIHTGTTYYRPSLESRLTIAMDPSKNQVSLSLTSVTVADSAMYYCVRIRGGSSNWLDPWGPGIVVTASS

#### Wildtype light chain

QSVWTQPPSVSAAPGQKVTISCSGDDSILRSAFVSWYQQVPGSAPKLVIFDDRQRPSGIPARFSGSNSGTTATLDIAGLQRGDEADYYCAAWNGRLSAFVFGSGTKL

#### Germline heavy chain

QVQLQESGPGLVKPSETLSLTCTVSGYSISSGYYWGWIRQPPGKGLEWIGSIYHSGSTYYNPSLKSRVTISVDTSKNQFSLKLSSVTAADTAVYYCARIRGGSSNWFDPWGQGTLVTVSS

#### Germline light chain

QSVLTQPPSVSAAPGQKVTISCSGSSSNIGNNYVSWYQQLPGTAPKLLIYDNNKRPSGIPDRFSGSKSGTSATLGITGLQTGDEADYYCGTWDSSLSAFVFGTGTKV

#### CII-Cit-13 peptide (GEEGK-Cit-GARG)

NCC(=O)N<u>C@@</u>(CCC(=O)O)C(=O)N<u>C@@</u>(CCC(=O)O)C(=O)NCC(=O)N<u>C@@</u>(CCCCN)C(=O)N<u>C@@</u>(CCCNC(N)=O)C(=O)NCC(=O)N<u>C@@</u>(C)C(=O)N<u>C@@</u>(CCCNC(=N)N)C(=O)NCC(=O)O

#### A1At-HCit peptide (EEAPLKLSKAVH-HCit-AVLTIDEKGTEAAGA)

N<u>C@@</u>(CCC(=O)O)C(=O)N<u>C@@</u>(CCC(=O)O)C(=O)N<u>C@@</u>(C)C(=O)N1<u>C@@</u>(CCC1)C(=O)N<u>C@@</u>(CC(C)C)C(=O)N<u>C@@</u>(CCCCN)C(=O)N<u>C@@</u>(CC(C)C)C(=O)N<u>C@@</u>(CO)C(=O)N<u>C@@</u>(CCCCNC(N)=O)C(=O)N<u>C@@</u>(C)C(=O)N<u>C@@</u>(C(C)C)C(=O)N<u>C@@</u>(CC1=CN=C-N1)C(=O)N<u>C@@</u>(CCCCN)C(=O)N<u>C@@</u>(C)C(=O)N<u>C@@</u>(C(C)C)C(=O)N<u>C@@</u>(CC(C)C)C(=O)N<u>C@@</u>(<u>C@</u>(O)C)C(=O)N<u>C@@</u>(<u>C@</u>(CC)C)C(=O)N<u>C@@</u>(CC(=O)O)C(=O)N<u>C@@</u>(CCC(=O)O)C(=O)N<u>C@@</u>(CCCCN)C(=O)NCC(=O)N<u>C@@</u>(<u>C@</u>(O)C)C(=O)N<u>C@@</u>(CCC(=O)O)C(=O)N<u>C@@</u>(C)C(=O)N<u>C@@</u>(C)C(=O)NCC(=O)N<u>C@@</u>(C)C(=O)O

The reference crystal structure for validation tests is derived by truncating the 5OCX PDB entry of E4 – CII-Cit-13 to only the variable domain of the antibody ^38^.

Predictions using various structure prediction methods were performed as follows: Chai-1: Predictions are conducted with multiple sequence alignments (MSAs) enabled, no template, no specific restraints, and default flags (3 trunk recycles and 200 diffusion timesteps) ^40^. AlphaFold3: Predictions are made with default settings (10 recycles, 1 diffusion sample, and 1 seed) ^41^. Boltz-1: Predictions are completed using MSA mode “mmseqs2_uniref_env,” with 5 recycles, 200 sampling steps, and 1 diffusion sample ^44^. RoseTTAFold All-Atom: Predictions are executed with MSA search enabled ^42^. Protenix: Predictions are performed with MSA search and ESM features ^43^.

### Calculation of Root Mean Square Deviations (RMSD) for CDR Loops Compared to Crystal Structures

To assess the accuracy of the predicted complementarity determining region (CDR) loops, we calculate the root mean square deviations (RMSD) between the predicted structures and the corresponding crystal structure.

The RMSD for CDR loops and the framework region is calculated using the PyRosetta library ^63^.

The command to execute the RMSD calculation is as follows:

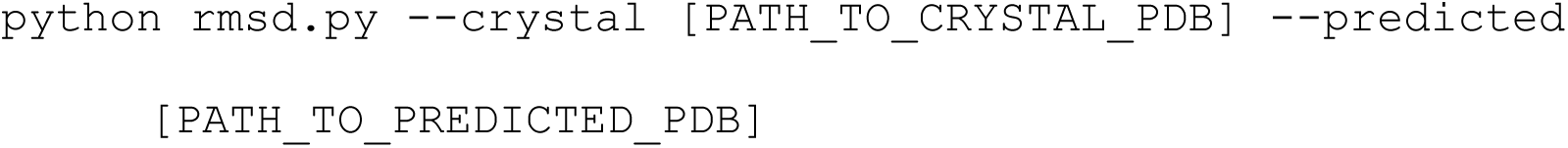

In this command:

`--crystal` specifies the path to the crystal structure PDB file.

`--predicted` specifies the path to the predicted structure PDB file.

### Calculation of Global Biophysical Properties

Global biophysical properties of the antibodies E4 and E4GL, including net charge, paratope charge, paratope solvent accessible surface area (SASA), and VH/VL packing angles and distances, are calculated using the Rosetta AntibodyFeatures reporter ^64^.

To execute the calculation, use the following command:

~~~
python AntibodyFeatures.py [PATH_TO_UNBOUND_PDB]
~~~

### Calculation of Per-CDR Biophysical Properties

Per-CDR biophysical properties of E4 and E4GL are derived by querying the SQL database generated from the AntibodyFeatures reporter.

To perform this analysis, run the following command:

~~~
python db3_check.py --db3 [PATH_TO_DB3]
~~~

### Calculation of Binding Affinity for Antibody-Peptide Complexes

Binding affinity between antibody-peptide complexes is predicted using PRODIGY-LIG, a structure-based method designed for the prediction of binding affinity in protein-small ligand complexes ^65^.

The binding affinity is calculated using the following command:

~~~
prodigy_lig -c A,B,C C:LIG -i [PATH_TO_BOUND_PDB]
~~~

In this command:

`--c A,B,C` specifies the chains of the antibody involved in the interaction

`C:LIG` indicates that the ligand (peptide) is associated with chain C

`--i [PATH_TO_BOUND_PDB]` is the path to the PDB file containing the antibody-peptide complex. The default distance cutoff for atomic contacts is set to 10.5 Å, which has been established as optimal for these calculations.

### Identification of Intermolecular Interactions

Intermolecular interactions between the antibody and the peptide are visualized using PyMol. Specifically, the citrulline (or homocitrulline) within the peptide is selected, and the following command identifies any polar contacts between the residue and the heavy or light chain of the antibody:

~~~
actions > find > polar contacts > to any atoms
~~~

## Author contributions

Conception: FA. Designing research studies: MES, MC, JJG, FA. Conducting experiments: MES, MC, DF, JJG, FA. Data acquisition and analyses: MES, MC, DF, JJG, FA. Interpretation of data: MES, MC, DF, JJG, FA. Writing—original draft: MES, MC, JJG, FA. Writing—review and editing: MES, MC, DF, JJG, FA. All authors reviewed, edited, and approved the manuscript.

## Data and materials availability

All data are available in the main text or the supplementary materials.

