## Supplemental for "Anti-citrullinated protein antibodies arise during affinity maturation of germline antibodies to carbamylated proteins in rheumatoid arthritis"

**This file contains:**

Supplementary Figs. 1-3
Supplementary Tables 1-4

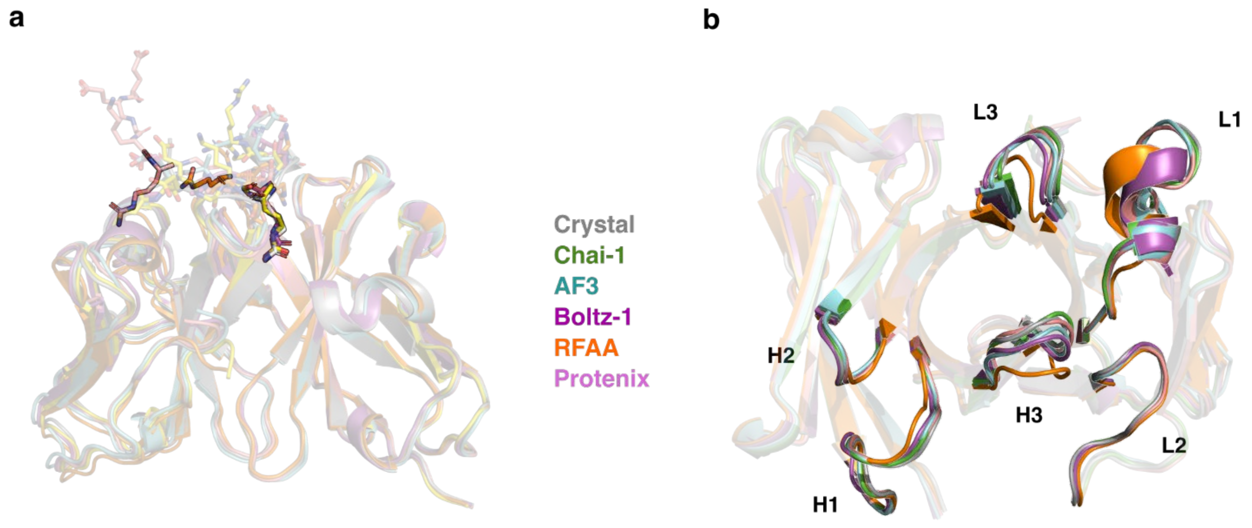


**Supplementary Fig. 1. Validation of structure prediction method.** A set of AI methods for biomolecular structure prediction are benchmarked against a known crystal structure of E4 with a citrullinated CII-Cit-13 peptide (PDB: 5OCX, gray). Each of the predictive models are derived by providing an amino acid sequence of E4 variable domain sequence and a SMILE sequence of CII-Cit-13 peptide. Comparison of (**a**) CII-Cit-13 conformation and (**b**) predicted CDR loop structures for target 5OCX for Chai-1 (green), AlphaFold3 (cyan), Boltz-1 (cyan), RoseTTAFold-AA (orange), and Protenix (pink) versus the experimentally determined structure (gray). The results support that Chai-1 is best for predicting the structure of E4GL and its bound conformation with citrullinated and carbamylated peptides.

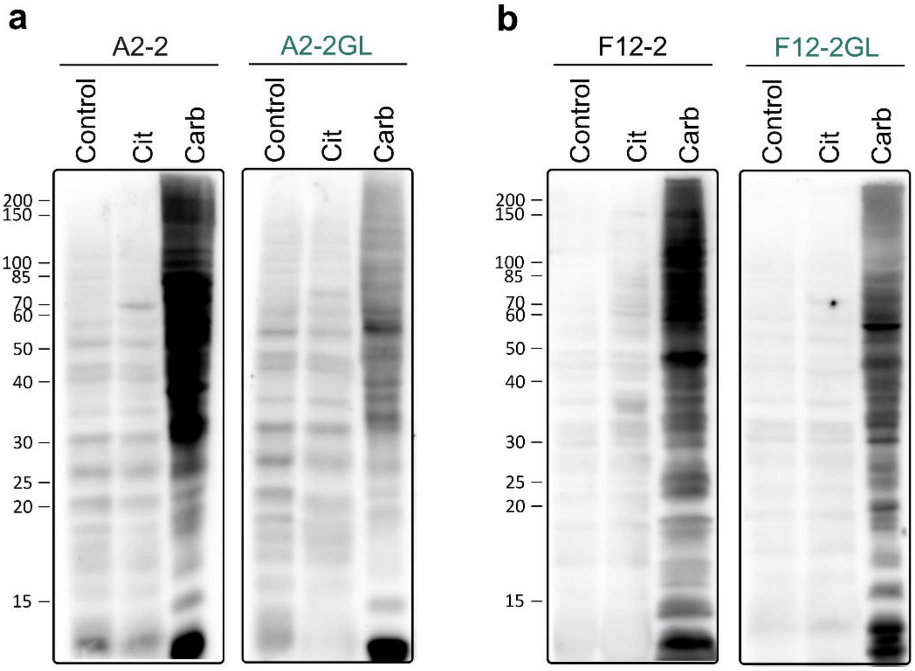


**Supplementary Fig. 2. RA-derived monoclonal antibodies A2-2 and F12-2, as well as their germline reverted forms, primarily recognize carbamylated proteins.** (**a, b**) Lysates from control, citrullinated (Cit), and carbamylated (Carb) cells were tested by immunoblotting using 6.6 nM of antibodies A2-2, A2-2GL (**a**), F12-2 or F12GL (**b**). Representative data from two independent experiments are shown.

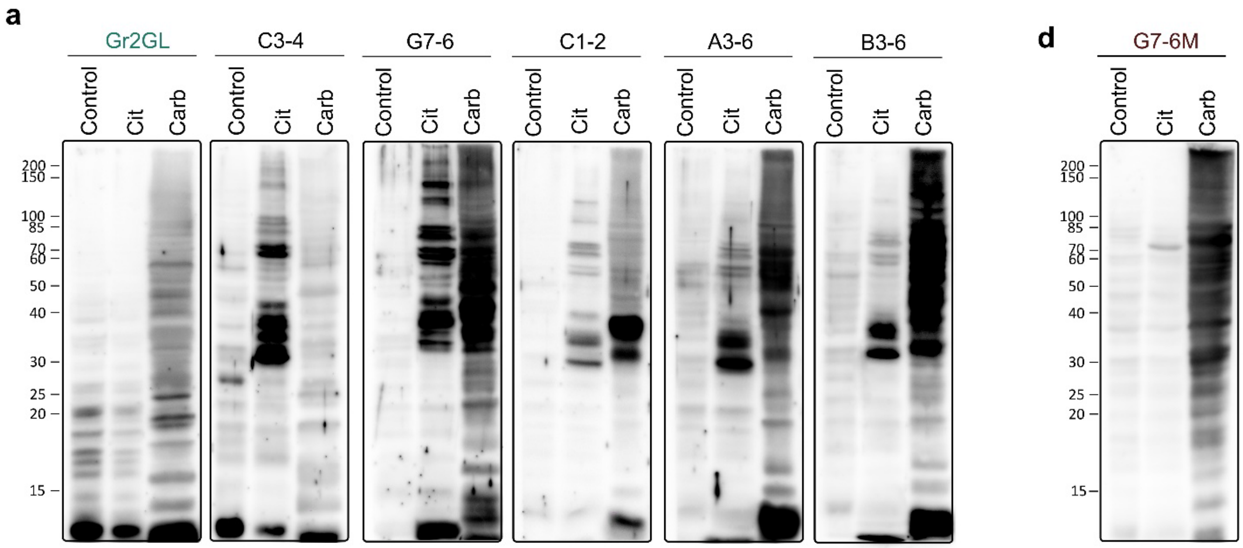


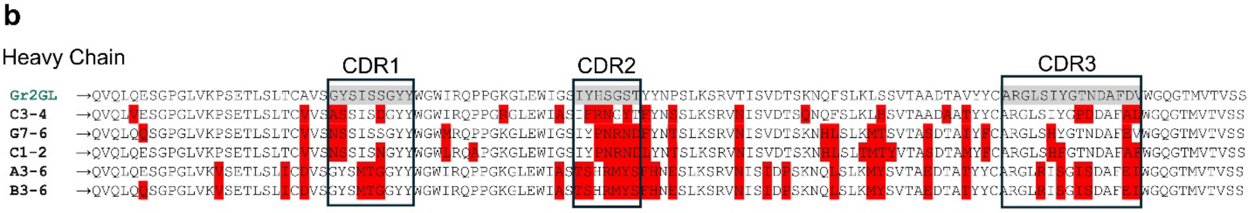


**c**

Missense Mutations

| | TOTAL | FR1 | CDR1 | FR2 | CDR2 | FR3 | CDR3 | FR4 |
|---|---|---|---|---|---|---|---|---|
| Gr2GL | 0 | 0 | 0 | 0 | 0 | | 0 | 0 |
| C3-4 | 22 | 2 | 3 | 2 | 4 | 7 | 4 | 0 |
| G7-6 | 22 | 2 | 2 | 1 | 5 | 10 | 2 | 0 |
| C1-2 | 27 | 1 | 3 | 2 | 5 | 12 | 4 | 0 |
| A3-6 | 30 | 3 | 3 | 1 | 6 | 11 | 6 | 0 |
| B3-6 | 31 | 4 | 3 | 1 | 6 | 11 | 6 | 0 |

**Supplementary Fig. 3. A single germline-encoded anti-CarP antibody generates anti-CarP, ACPAs, and dual-reactive anti-CarP/ACPAs as result of somatic hypermutations.** (**a**) Lysates from control, hypercitrullinated (Cit) and carbamylated (Carb) cells were tested by immunoblotting using antibody Gr2GL, C3-4, G7-6, C1-2, A3-6 or B3-6. (**b, c**) VH amino acid sequence alignment (**b**) and number of missense mutations (**c**) of antibodies shown in **a**. In **b**, missense mutations are marked in red. FR, framework region. CDR, complementarity-determining region. (**d**) Lysates from control, Cit and Carb cells were tested by immunoblotting using antibody G7-6M. All RA-derived monoclonal antibodies were tested at a concentration of 6.6 nM. Data in **a** and **d** are representative of 3 independent experiments.

**Supplementary Table 1. Benchmarking structure prediction models.** Performance of various biomolecular structure prediction models at predicting the structure of E4. Numbers are the root mean squared deviations in angstroms of each predicted antibody structural segment relative to the experimental structure after superimposing the entire Fv region ($V_H$ and $V_L$ domains).

| | RMSD | | | | | | | |
|---|---|---|---|---|---|---|---|---|
| | H1 | H2 | H3 | FR H | L1 | L2 | L3 | FR L |
| **Chai-1** | 0.30 | 0.20 | **0.36** | **0.26** | 0.45 | 0.19 | 0.35 | **0.28** |
| **AF3** | 0.30 | **0.16** | 0.63 | 0.28 | 0.74 | **0.16** | **0.20** | 0.45 |
| **Boltz-1** | 0.54 | 0.19 | 0.48 | 0.43 | 1.38 | 0.18 | 0.83 | 0.36 |
| **RF-AA** | 0.32 | 0.67 | 0.41 | 0.72 | 0.44 | 0.20 | 0.60 | 1.10 |
| **Protenix** | **0.23** | 0.32 | 0.38 | 0.32 | **0.43** | **0.16** | 0.69 | 0.32 |

**Supplementary Table 2. In-silico predicted biophysical properties of E4 and E4GL variable fragment.** Biophysical properties are predicted with Rosetta's AntibodyFeatures [64] object given the structure of E4-CII-Cit-13 and E4GL-A1AT-HCit predicted from Chai-1 as input. SASA is solvent accessible surface area.

| | **E4** | **E4GL** |
|---|---|---|
| Net charge ($e^-$) | 0 | 4 |
| Paratope charge ($e^-$) | 1 | 2 |
| Paratope SASA ($Å^2$ ) | 2685.4 | 2239.5 |
| Paratope hSASA ($Å^2$) | 1161.9 | 903.6 |
| Paratope pSASA ($Å^2$) | 1523.5 | 1336.0 |
| VL/VH packing angle (°) | 89.2 | 89.8 |
| VL/VH distance (Å) | 17.5 | 17.6 |
| VL/VH opening angle (°) | -35.4 | -36.3 |
| VL/VH opposite opening angle (°) | 64.5 | 66.9 |

**Supplementary Table 3. In-silico predicted biophysical properties of E4 and E4GL CDR loops.** Biophysical properties are predicted with Rosetta AntibodyFeatures [64] given the structure of E4-CII-Cit-13 and E4GL-A1AT-HCit complexes predicted from Chai-1 as input. SASA is solvent accessible surface area.

| | E4 | | | E4GL | | |
|---|---|---|---|---|---|---|
| | SASA (Å$^2$) | Charge ($e^-$) | Aromatic Residues | SASA (Å$^2$) | Charge ($e^-$) | Aromatic Residues |
| **H1** | 630.8 | -1 | 4 | 519.0 | 0 | 4 |
| **H2** | 359.7 | 0 | 0 | 350.2 | 0 | 1 |
| **H3** | 262.7 | 2 | 2 | 255.2 | 2 | 2 |
| **L1** | 606.8 | -1 | 1 | 414.8 | 0 | 1 |
| **L2** | 451.7 | 0 | 1 | 380.0 | 1 | 1 |
| **L3** | 373.7 | 1 | 1 | 320.3 | -1 | 1 |

**Supplementary Table 4. In-silico predicted binding affinities of antibody-peptide complexes**. Binding affinity between the antibody and the non-canonical amino acid (citrulline or homocitrulline) within the peptide are calculated using PRODIGY-LIG [65].

| Complex | Binding affinity (ΔG Kcal/mol) |
|---|---|
| E4 – Cit (from CII-Cit-13) | -6.01 |
| E4 – HCit (from A1AT-HCit) | -5.41 |
| E4GL – Cit (from CII-Cit-13) | -5.80 |
| E4GL – HCit (from A1AT-HCit) | -5.93 |
